# Streamlining Asymmetry Quantification in Fetal Mouse Imaging: A Semi-Automated Pipeline Supported by Expert Guidance

**DOI:** 10.1101/2024.10.31.621187

**Authors:** S.M. Rolfe, D. Mao, A. M. Maga

## Abstract

Asymmetry is a key feature of numerous developmental disorders and in phenotypic screens is often used as a readout for environmental or genetic perturbations to normal development. A better understanding of the genetic basis of asymmetry and its relationship to disease susceptibility will help unravel the complex genetic and environmental factors and their interactions that increase risk in a range of developmental disorders. Large-scale imaging datasets offer opportunities to work with sample sizes needed to detect and quantify differences in morphology beyond severe deformities while also posing challenges to manual phenotyping protocols. In this work, we introduce a semi-automated open-source workflow to quantify abnormal asymmetry of craniofacial structures that integrates expert anatomical knowledge. We apply this workflow to explore the role of genes contributing to abnormal asymmetry by deep phenotyping 3D fetal microCT images from knockout strains acquired as part of the Knockout Mouse Phenotyping Program (KOMP2). Four knockout strains: *Ccdc186*, *Acvr2a*, *Nhlh1*, and *Fam20c* were identified with highly significant asymmetry in craniofacial regions, making them good candidates for further analysis into their potential roles in asymmetry and developmental disorders.

**Summary Statement:** We introduce an open-source, semi-automated pipeline to detect abnormally asymmetric phenotypes in 3D scans of fetal mice to explore the relationship between facial asymmetry, perturbed development, and developmental instability.

## Introduction

Asymmetry is a trait frequently associated with disorders such as schizophrenia, autism spectrum disorder, fetal alcohol exposure disorders and a number of structural birth defects such as orofacial clefting, microtia, Cornelia de Lange syndrome, in addition to many others (Naugler et al., 1996; Gourion et al. 2004; Weinberg et al., 2006; Luquetti et al., 2012; Mehta et al., 2016; Postema et al., 2019; Blanck-Lubarsch et al., 2020). As construction of symmetric structures is controlled by the same set of genes, differences outside of reflection or rotation can be evidence of either a breakdown in regulation, or the presence of a control process impacting expression (Naugler et al., 1996; Boorman et al., 2002; Graham et al., 2021). Two types of asymmetries can be distinguished in a population by the distribution of differences between the left and right sides of the plane of symmetry: fluctuating asymmetry (FA) and directional asymmetry (DA). FA is defined as random bilateral deviations from normal symmetry and has long been used as a measure of developmental instability (DI), due either to genetic or environmental causes (Mather et al. 1953; Ekrami et al., 2018; Varón-González et al., 2019). Studies have linked FA to a range of disorders including major, structural birth defects such as oral facial clefts (OFCs) and craniofacial microsomia (CFM) and neurological disorders including schizophrenia, autism spectrum disorder, and Down’s syndrome (Starbuck et al., 2013; Stephan-Otto et al., 2020). Directional asymmetry (DA) is defined as asymmetry in which one side of a bilateral structure is consistently larger than the other. While DA can be a normal property of an organism (e.g., consistent differences in left and right hemispheres of brain), like FA, DA is also associated with developmental disorders, including OFC and CFM (Hammond et al., 2008). Evidence for involvement of DA in OFC comes from both the clinical presentation of clefts, which is often unilateral, with a higher rate of incidence on the left side, and high rates of craniofacial DA measured in unaffected parents of children with cleft lip (Zemann et al., 2002; McIntyre et al., 2010). Specifically, craniofacial asymmetry measured in relatives was shown to be predictive of the side of unilateral clefting in affected individuals (Yoon et al., 2003). Links between FA, DA, and a range of developmental disorders suggests that genetic and environmental factors may be shared or operate synergistically (Neiswanger et al., 2002; Neiswanger et al., 2005). Finding novel variants contributing to asymmetry and assessing their possible role in developmental disorders with an asymmetric phenotype will help elucidate the complex etiology of these disorders and have the potential to improve preventive measures, diagnostics and therapies.

The International Mouse Phenotyping Consortium (IMPC) has led a systematic phenotype evaluation of targeted knockout mouse strains with the goal of providing a comprehensive dataset for exploring genotype/phenotype interactions through projects like Knockout Mouse Phenotyping Project 2 (KOMP2), an international project that includes 18 institutes and centers (Wong et al., 2012; Dickinson et al., 2016). To date, 7,022 of the 23,000 targeted genes have been evaluated. As part of the standardized phenotyping pipeline, high-resolution 3D microCT fetal imaging is collected for sub-viable and lethal strains. As of 2024, there are 267 lines at E14.5/E15.5 with an average of 6 embryos. The high-resolution microCT imaging data, along with a large wild-type control group provides the capability to detect subtle differences in size and shape, such as mild asymmetry of individual structures or organs with the right approach. Since these knockout lines are sub-viable or embryonically lethal, the gene whose function suppressed likely represent a major developmental gene or pathway crucial for the normal development. Analysis of morphological differences in these knockout strains compared to the baseline specimens presents an opportunity to test the hypothesis of association of increased asymmetry with likelihood of structural abnormality.

Large scale imaging datasets such as provided by KOMP2 are becoming increasingly prevalent and unlock the potential for detecting nuanced phenotypes with greater statistical significance. However, when dealing with datasets of sufficient size to detect smaller scale phenotypic abnormalities, manual assessment of each image by human experts is simply not feasible. This challenge is especially pronounced when the phenotypes are subtle or difficult to identify and require careful quantification. Systematic methods to automate quantification of developmental abnormalities are necessary for projects on this scale to leverage the rich image information captured by high resolution scanning.

Quantifying abnormal asymmetry from a 3D image is a complicated task. Differences in pose, alignment, and the presence of normally occurring asymmetry complicate visual assessment. Standard phenotyping protocols using an extensive set of landmark points placed at repeatable locations across an image can provide a quantification of differences between bilateral sides, however repeatable points on a structure such as an embryo are very sparse, more so in the cases of significantly malformed embryos. Surfaces that are of high biological importance with curved or smooth surface geometry, such as the maxillary and mandibular regions, may have very few identifiable points (i.e., distinct anatomical landmarks) available. This can result in a sparse representation of shape that is not capable of describing asymmetries in surface curvature or orientation in these regions.

Another reality of dealing with large scale imaging datasets, often collected from multiple imaging centers, is the variation in the image quality. Scans may exhibit imaging artifacts or show post-mortem damage to the specimens that might have occurred during the preparation of the specimen for imaging. Detecting aberrations in images and separating them from developmental changes to morphology is a crucial task of automated analysis techniques. While it may be ideal to exclude specimens with soft tissue damage, the small sample sizes of knockout strains or other difficult to obtain experimentally manipulated groups was the primary motivation in our development of techniques that are robust to this type of noise outside the region of interest.

In this study, we demonstrate a semi-automated, expert guided, fully open-source approach to spatially dense quantification of abnormal asymmetry applied to analyze facial asymmetry in the KOMP2 3D embryonic microCT scan data with the goal of identifying knockout strains with heightened asymmetry.

## Methods

### Alignment landmarking

The initial step in our asymmetry quantification pipeline is the manual annotation of landmarks. The 17 landmarks collected for each specimen are described in Figure 1. These landmarks are divided into two sets, global positioning landmarks used to align the specimen in a common orientation for analysis (landmarks 1-14) and local landmarks that provide anatomical homology between specimens in regions of high shape variability that assist in determining the point correspondences during the deformable stage of the mapping (landmarks 15-17). The choice of these landmark sets is a key design step when using the pipeline and is dependent on the image geometry.

**Figure 1.**
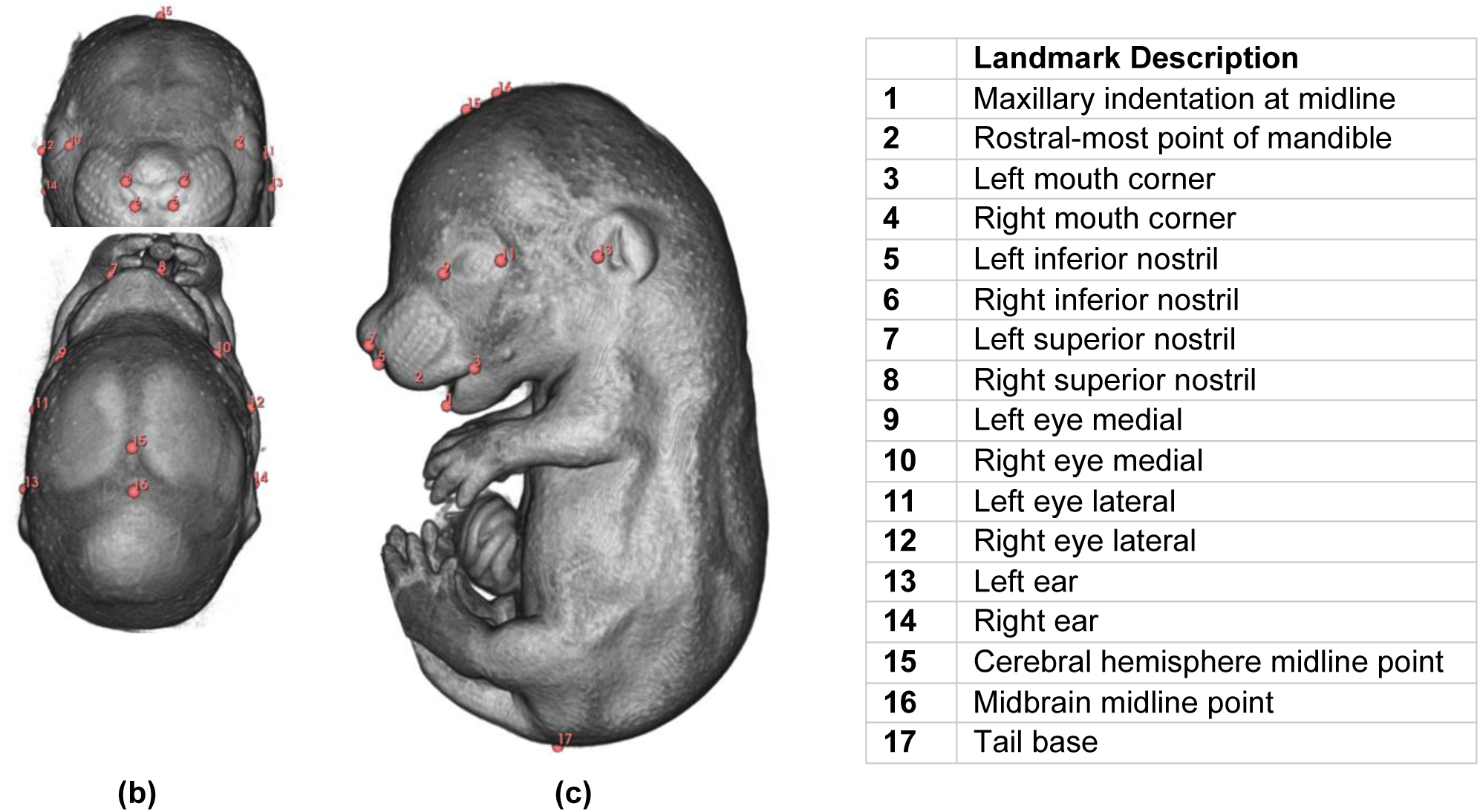
17 manually annotated landmark definitions. Landmarks 1 through 14 are used for global positioning. Landmarks 15 through 17 are included to capture local shape differences to improve the accuracy of point correspondence assignment.

To assess facial asymmetry, 14 global alignment landmarks were chosen from the craniofacial region. These global landmarks are used for a rigid alignment that minimizes the Euclidean distance between each individual landmark configuration and the atlas, distributing the difference between the landmark points. Including a landmark from a known point of high variation at this step will result in larger errors in the remaining alignment landmarks. Plastic deformation of the limbs and body curvature represent the largest source of variation in this dataset. Accounting for these local shape and positional differences is important in the deformable step for establishing accurate point correspondences but they should be excluded from the global rigid registration, as they will introduce error into the alignment of the facial region. Three local landmark points were selected from the cranial and tail regions to capture shape difference in the body in the deformable registration step of this algorithm, where local landmarks are combined with the global positioning landmarks as the basis for a thin plate spline (TPS) transformation that removes shape differences prior to the assignment of point correspondences.

While the manual landmarking can be tedious when done from the scratch as we noted previously, tools provided in the open-source 3D Slicer image analytics platform (Fedorov et al., 2012; Kikinis et al., 2014) and its SlicerMorph extension (Rolfe et al, 2023b) make it more streamlined. We have specifically made use of the volume rendering presets that provided a consistent 3D rendering of all embryos (diceCT preset), and a blank landmark-template, which made sure the 17 landmarks were always recorded in the same order, using the same visualization style. Using these two features, all 17 landmarks from a single embryo can be recorded in less than a minute by someone familiar with anatomy.

### Extracting embryo surface for analysis

The first step in the analysis pipeline is the segmentation of the external surface of embryos. The goal of this task is to split the image into a single foreground and background segment. Manual segmentation of a large dataset is time-consuming and poses hurdles to further quantitative shape analysis. However, simple automated solutions such as applying a threshold computed from image properties are often not effective enough due to variability in imaging parameters, noise in the image, and internal structures created within the image surface. To improve the efficiency of the analysis pipeline, we have trained a deep learning model that can be run within the 3D Slicer application to assist with the extraction of the embryo surface. Deep learning networks have recently gained attention as a competitive alternative with the potential to improve performance over automated methods requirements (Hatamizadeh, 2022; Schoppe, 2020; Tang, 2021). This is especially beneficial when the anatomy is highly varied and may not be well represented by a reference segmentation as it is required by atlas-based methods or may require significant user interaction if segmented fully manually. While training a deep learning model is a computationally intensive task, a fully trained network is comparatively lightweight and can estimate a segmentation in a fraction of the time (Rolfe et al, 2023a). The deep learning segmentation tool we have produced for this work is planned to be made available as part of the MEMOS extension to the 3D Slicer platform, providing a quick and convenient method to estimate a segmentation that can be reviewed, and if necessary edited within the same application as part of this analysis workflow.

#### Model training

The model we trained for whole-body segmentation utilizes a UNet with Transformers (UNETR) architecture as described in Hatamizadeh et. al. (2022) with encoding and decoding units arranged in a UNet-like contracting-expanding pattern, implemented using the MONAI library. The training data was generated by manual segmentation of the embryo surface from a set of 91 baseline scans. The labeled dataset was partitioned into a set of 82 training scans and 20 test scans. The model was trained for 14,000 iterations on a server with 512 GB RAM and A6000 GPUs.

An optional automated post-processing step is included in the surface extraction pipeline to fill any internal holes in surface segmentation, creating a closed surface. While our model was trained for segmentation of KOMP2 acquired data, this segmentation model can be tuned to create customized segmentation models for specific datasets by additional training with E15.5 fetal mouse scans from other sources, such as microCT scans acquired at institutions using different scan settings or imaging protocols.

### Symmetry analysis

The symmetry assessment workflow was developed as a part of our group’s 3D Slicer extension, DeCA: Dense Correspondence Analysis Toolkit (Rolfe et al. 2023b). DeCA employs a semi-automated approach to map corresponding points across surface models to a symmetric reference atlas. User-provided landmarks serve as anchors for automatically assigned point correspondences based on geometric similarity. Asymmetry is quantified for each model by comparing the position of bilaterally symmetric points. This method has previously been used by our group to quantify facial asymmetry in surface scans and was adapted for this application to allow for greater anatomical variation (Rolfe et al., 2018). The DeCA module, implemented in the open-source 3D Slicer application, provides an interface that guides the user through each step of the symmetry assessment and is publicly available through GitHub (https://github.com/smrolfe/DeCA). DeCA was used to build a computational model of normal asymmetry for the control group and quantify pointwise asymmetry in the KO strains through the steps outlined below.

#### Atlas creation

A symmetrized average model of normal anatomy for fetal stage E15.5 was generated from baseline images in the KOMP database. All baseline images are registered to the specimen lying closest to the Procrustes mean shape of the manual landmark points, using the two-step rigid and deformable transform. Corresponding point position for each vertex on the reference specimen model are averaged and symmetrized across the baseline dataset. The resulting model serves as an atlas of the point correspondences for asymmetry analysis.

#### Pointwise asymmetry

To calculate pointwise asymmetry associated with the midsagittal plane, we generated a mirror image of each specimen. Bilaterally symmetric landmarks points are mirrored and reordered according to their mirrored positions. The original image and the mirror image copy are registered to the atlas image. At each atlas vertex, asymmetry is represented by the magnitude of the vector between the corresponding points on the original and mirrored image mapped to that vertex, as shown in Figure 2.

**Figure 2.**
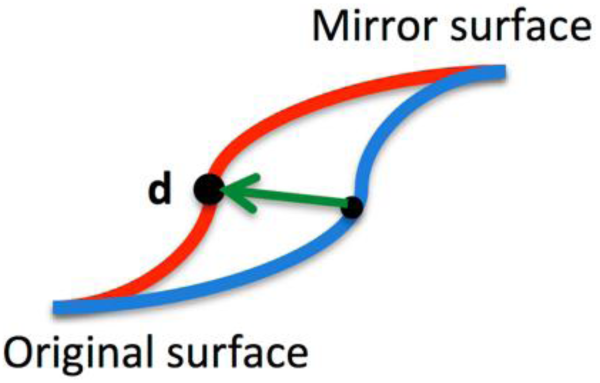
Pointwise calculation of asymmetry: Magnitude is calculated as the length of the vector *d* between points on the original and mirrored surfaces that are mapped to the same vertex point in the atlas.

#### Asymmetry scoring

To score craniofacial asymmetry, 5 regions of interest are selected on the atlas image: the nose, mandibular region, maxillary region, eyes and otic placode shown in Figure 3. These regions correspond to the 5 facial regions described in the FaceBase Atlas of C57BL/6 Mouse Embryo Anatomy (https://www.facebase.org/resources/mouse/mouseanatomy). The regions are segmented on the atlas model in the 3D Slicer application via curved lines placed on the image. Each facial region is assigned an asymmetry score by averaging the pointwise asymmetry at each vertex in the region. Since the regions are placed on the atlas image after asymmetry quantification, symmetry scores for new regions can be calculated without rerunning the analysis.

**Figure 3.**
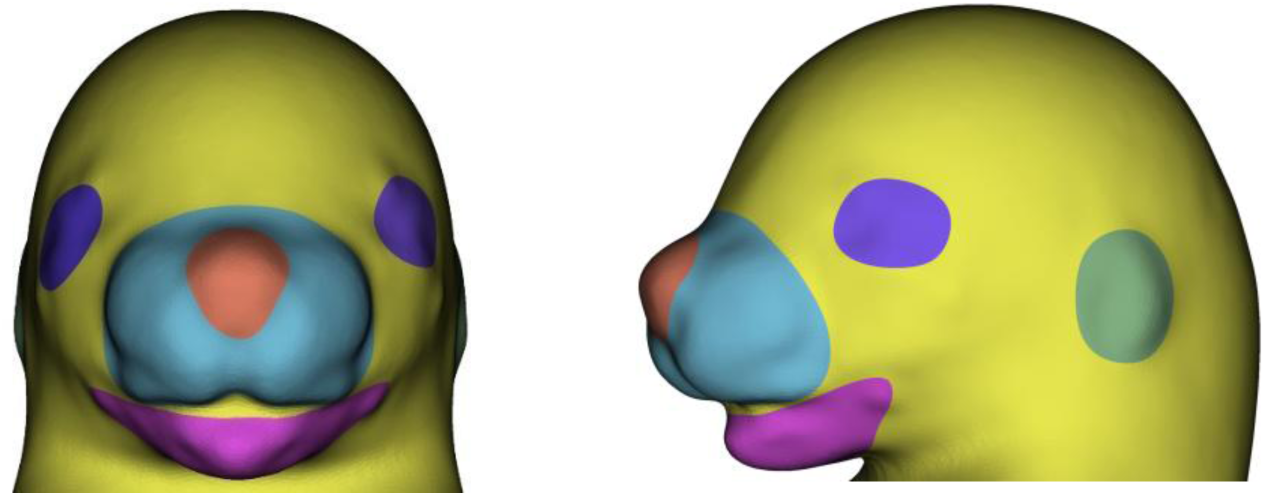
Facial subregions identified from FaceBase Atlas of C57BL/6 Mouse Embryo Anatomy including the nose, mandibular region, maxillary region, eyes and otic placode.

## Results

The model of normal facial asymmetry was generated from a group of 95 baseline specimens from the KOMP2 dataset. A symmetric average atlas was created and symmetrized as an index of the corresponding points assigned across the dataset. Each baseline specimen and its mirrored copy were warped to the baseline atlas and the magnitude of asymmetry was calculated at each point. The average magnitude of asymmetry displayed as a heatmap on the baseline atlas is shown in Figure 4.

**Figure 4.**
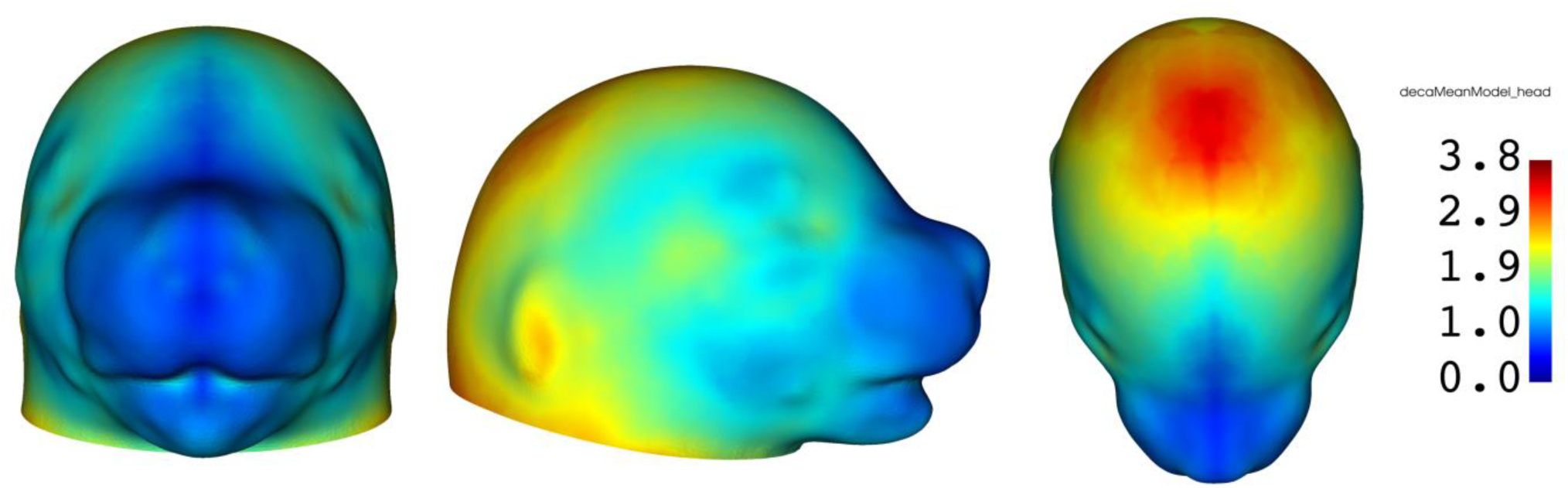
Mean magnitude of pointwise asymmetry for baseline strains displayed as a heatmap where red indicates the highest magnitude and blue represents the lowest magnitude differences.

Specimens from 53 KO strains were screened for post-mortem damage, image artifacts, or gross structural malformations in the craniofacial region. Specimens were excluded from further analysis when these factors prevented acquisition of a reliable facial surface for analysis or identification of anatomical correspondences. After this initial screen, asymmetry in the craniofacial region was quantified for 168 specimens from 31 KO strains and 95 specimens from the baseline strain. The KO strains and their sample sizes are detailed in the Supplementary Table S1.

### Significance of craniofacial asymmetry in perturbed development

To assess the relationship between craniofacial asymmetry and genetic perturbations to normal development, we pooled 133 specimens from 23 KO strains with no recorded structural phenotype in the KOMP2 database and compared the asymmetry scores from the 5 craniofacial regions to those from the 95 baseline specimens. These strains are listed in the Supplementary Table S2. As shown in Table 1, statistically significant asymmetry in the KO group was found for the nose, maxillary region and otic placode, supporting the hypothesis linking perturbed development and asymmetric phenotypes.

**Table 1:** Significance of asymmetry from KO strains with no structural phenotype recorded in the KOMP database. Regions with P values higher than 0.05 were denoted as N/S.

| Nose | Maxillary region | Mandibular region | Eyes | Otic Placode |
| --- | --- | --- | --- | --- |
| p<0.01 | p<0.05 | N/S | N/S | p<0.05 |

### Identifying KO strains with asymmetric phenotypes

Specimens with the highest craniofacial asymmetry scores were used to prioritize KO strains for further analysis. 168 specimens from 31 KO strains were evaluated and ranked by the magnitude of total facial asymmetry. The most asymmetric specimens from the top 4 KO strains were: *Ccdc186, Acvr2a*, and *Nhlh1* and *Fam20c*. The statistical significance of the scores from each region for these specimens is detailed in Table 2 and images of the craniofacial region from each specimen is provided in the Supplementary Figure S1. The number of specimens with statistically significant asymmetry for each of the knockout strains in Table 2 is provided in Supplementary Table S3.

**Table 2:** Significance of asymmetry scores from four specimens with the highest asymmetry. Regions with P values higher than 0.05 were denoted as N/ S.

| Strain | Nose | Maxillary region | Mandibular region | Eyes | Otic Placode |
| --- | --- | --- | --- | --- | --- |
| *Ccdc186* | N/S | p<0.01 | p<0.01 | p<0.01 | p<0.01 |
| *Acvr2a* | p<0.05 | p<0.01 | N/S | p<0.01 | p<0.01 |
| *Nhlh1* | p<0.01 | N/S | N/S | p<0.01 | p<0.01 |
| *Fam20c* | p<0.01 | p<0.01 | p<0.01 | p<0.01 | N/S |

Within the KOMP2 phenotyping protocol, the knockout strains *Ccdc186* (2 out of 7 homozygotes screened) and *Acvr2a* (1/2 homozygotes screened) were associated with abnormal craniofacial morphology. *Nhlh1* and *Fam20c* were not reported as having a craniofacial phenotype. In KOMP2 pipeline, asymmetry is not evaluated as part of the phenotyping process and so may be present with or without other abnormalities in facial morphology.

Known links to genetic disorders that can disrupt facial development in humans may provide evidence of a gene’s involvement in pathways related to facial morphology. Mutations on three of the selected genes have known links to facial dysmorphology reported in the literature. In humans, mutations on CCDC186 are associated with abnormal head size and facial features (microcephaly, achondrogenesis) (Brugger et al., 2021). Mutations on ACVR2A are linked to craniofacial disorders including Multiple Synostosis Syndrome that has characteristic facial features including a broad, tubular-shaped nose and a thin upper vermilion (Monies et al., 2017, Timberlake et al., 2023). Mutations on FAM20C are associated in humans with Raine syndrome, characterized by craniofacial anomalies including microcephaly, low set ears, osteosclerosis, a cleft palate, gum hyperplasia, a hypoplastic nose, and eye proptosis (Simpson et al., 2011; Fradin et al., 2011). NHLH1 is not a well-studied gene and we found no published links to craniofacial anomalies.

Patterns of gene expression in humans also provide an opportunity to validate a gene’s relevance to facial morphology. The four candidate genes were evaluated using tissue expression data from the Genotype-Tissue Expression (GTEx) database. While the GTEx database provides only a breakdown of expression for tissue types, without information on anatomical location, we would expect a gene relevant to craniofacial development to include expression in tissue types found in the craniofacial region. Genes primarily expressed in tissue types not present in this region would be less motivating candidates for future study. CCDC186, ACVR2A, NHLH1 and FAM20C showed widespread expression across tissue types in humans, as shown in Figure 5 for CCDC186. The bulk tissue expression for ACVR2A, NHLH1 and FAM20C are included in the Supplementary Figure S2.

**Figure 5.**
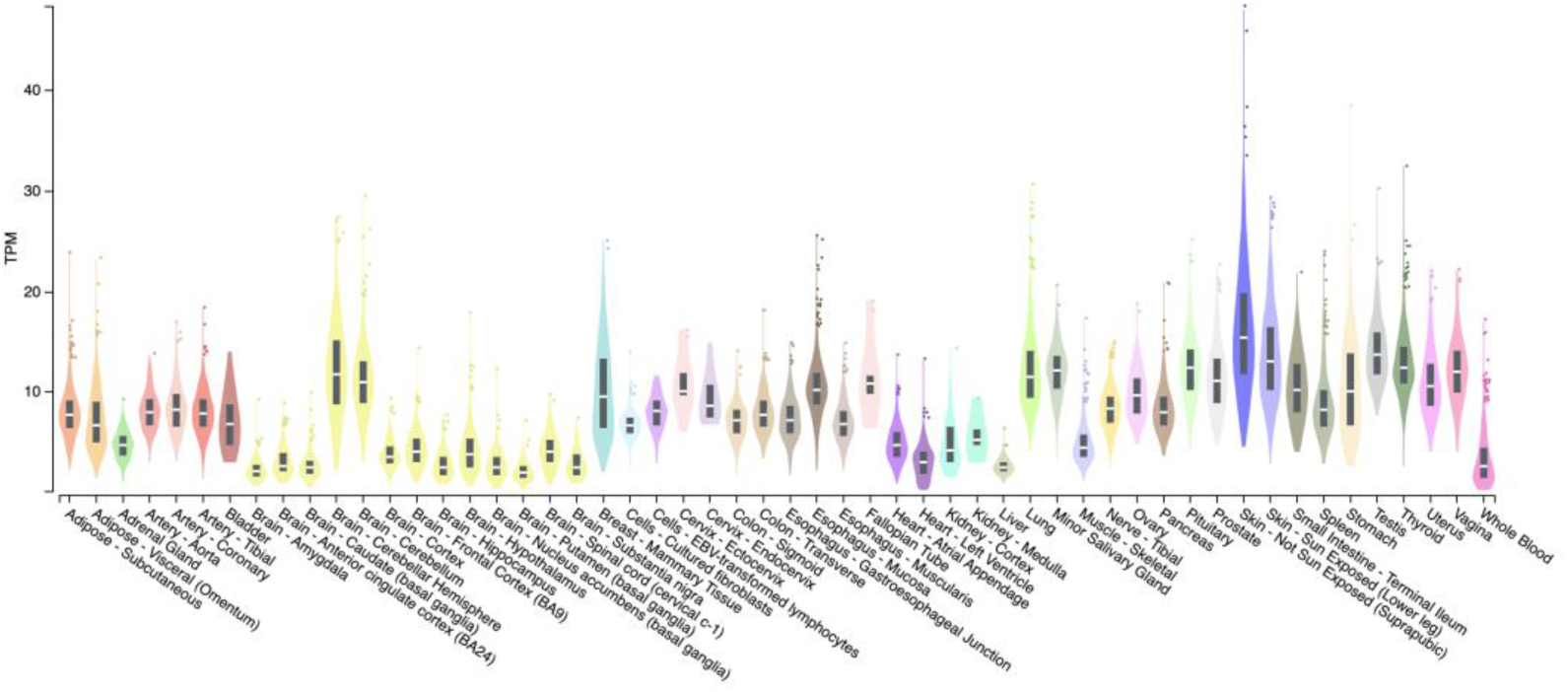
Bulk tissue gene expression from the GTEx database for *CCDC186*, showing widespread expression of the gene across sampled tissue types in humans.

## Discussion

In the KOMP2 dataset there are an unknown number of specimens that have soft tissue damage or imaging artifacts that would preclude them from shape analysis. Since the sample sizes for some knockout strain can be as low as 2-4 individuals, excluding damaged specimens can greatly reduce the power of the statistical analysis to detect differences. In this work we have assigned asymmetry scores based on craniofacial regions of interest identified on the atlas image. Restricting the quantification to specific regions of interest allows us to include damaged specimens when the tissue damage is outside these regions and provide local summaries of where significant asymmetry is occurring. Creating scores for new regions can be quickly done by simply drawing a curve on the full body atlas model in the 3D Slicer application that outlines the new region to be scored and does not require re-registering the sample to the atlas.

While our analysis pipeline was designed to provide a detailed quantification of asymmetry in regions of interest, it may also be used to flag highly abnormal images for manual review. Using whole body or whole head asymmetry is an effective way to screen for large magnitude differences that may indicate tissue damage, imaging artifact, or very severe anatomical differences such as missing limbs or exencephaly.

The most time-consuming step of the current pipeline is the manual landmarking step. However, this is a critical way to provide expert anatomical knowledge for the point correspondence assignment. While our landmarking and analysis workflow in 3D Slicer provides a highly streamlined process, in the future, automating the placement of these landmark points via deep learning models could be a route to improving efficiency.

### Separating fluctuating and directional asymmetry

The total magnitude of asymmetry between corresponding bilateral points in an individual can be broken down into directional asymmetry (DA), or the average asymmetry of a population, and fluctuating asymmetry (FA), measured as the random perturbations in an individual. While both DA and FA have links to developmental disorders, it has been argued that DA is attributable to genetic properties of a population while FA may be the result of genetic fitness or environmental stress and is typically used as a more direct proxy by which to measure DI (Zemann et al., 2002; Hammond et al., 2008; McIntyre et al., 2010; Yoon et al., 2003)

The standard way to calculate FA is to subtract the mean DA measured for the group from the total asymmetry of the individual. While this is approach is effective in population level studies, the DA component of asymmetry is not expected to be identical in each specimen. In an application where the goal is to screen individual specimens for a phenotype with heightened asymmetry, removing a constant, estimated DA component could mask the presence of FA. In this study, we opt to analyze the total magnitude of asymmetry, which does not distinguish between FA and DA, and report abnormality compared to the total magnitude of asymmetry measured in a control population.

## Conclusion

As 3D imaging becomes more available and widespread, there will be increasing opportunities to discover ‘deep’ phenotypes that capture and describe subtle differences in development, widening the range of phenotype/genotype associations that can be made. However, there are currently a large number of publicly available datasets that do not include this type of high - resolution phenotyping, and instead rely on dichotomous categories like ‘affected’ versus ‘unaffected’ labels, or other coarse measures.

This work demonstrates an open-source pipeline for high-density, deep phenotyping of 3D images and its potential to reveal genes implicated in abnormally asymmetric anatomy in the KOMP2 dataset. The analysis workflow provides an accessible and repeatable method for quantifying asymmetry and morphological abnormalities that requires no previous programming experience, purchase of equipment, or software. By automatically generating detailed descriptions of asymmetric phenotypes our tool provides a solution to the challenge of quantifying complex shape information from high-dimensional image data for large scale studies that may be applied in a range of research contexts

In our pilot analysis of the KOMP2 imaging data, we screened 53 KO strains and quantified asymmetry in 168 specimens from 31 KO strains and 95 specimens from the baseline strain. We prioritized 4 KO strains: CCDC186, ACVR2A, NHLH1 and FAM20C with statistically significant asymmetric craniofacial phenotypes for further investigation into the relationship between abnormal asymmetry and anomalous development. Hypothesis-driven selection of candidate genes for analysis, as we have shown in this work, is a powerful, complementary approach when working with high-dimensional whole genome sequencing data. It allows for a focus on variations in specific, biologically relevant regions of the genome with greater power, providing new opportunities to unravel the contributions of specific risk factors.

## Data Availability

The fetal mouse imaging data used in this work are publicly available through the International Mouse Phenotyping Consortium’s website at https://mousephenotyping.org. The source code for the DECA module, which can be run by manually installing as an extension of 3D Slicer, is available in the GitHub repository: https://github.com/smrolfe/DeCA.

The deep learning model for estimating surface contours from fetal mouse imaging and scripts used for training are available on request and are planned to be published as part of the SlicerMorph MEMOS extension to 3D Slicer: https://github.com/SlicerMorph/SlicerMEMOS.

## Acknowledgements

This work was supported by grants OD032627 and HD104435 to AMM from National Institutes of Health.

