## Supplemental Information for "Streamlining Asymmetry Quantification in Fetal Mouse Imaging: A Semi-Automated Pipeline Supported by Expert Guidance"

**Supplementary Information**

| Strain | Samples |
|---|---|
| *Acvr2a* | 5 |
| *Acvr2b* | 6 |
| *Aebp1* | 4 |
| *Aldh1a3* | 2 |
| *Aloxe3* | 4 |
| *Ampd1* | 7 |
| *Ano6* | 8 |
| *Arid1b* | 3 |
| *Baiap2* | 6 |
| *Bckdha* | 6 |
| *Casq2* | 8 |
| *Casr* | 4 |
| *Cbx4* | 3 |
| *Ccdc186* | 4 |
| *Cdkn1c* | 5 |
| *Chst14* | 7 |
| *Clps* | 8 |
| *Creb3* | 6 |
| *Dcc* | 8 |
| *Dok7* | 8 |
| *Fam20c* | 6 |
| *Hmox1* | 2 |
| *Krt5* | 6 |
| *Nabp2* | 5 |
| *Nhlh1* | 6 |
| *Pax7* | 4 |
| *Pdx1* | 1 |
| *Rfx6* | 6 |
| *Robo4* | 5 |
| *Slc6a5* | 5 |
| *Vars2* | 4 |
| *Wnt1* | 5 |

**Table S1: The sample sizes for the 31 KOMP2 knockout strains analyzed for asymmetry out of the 53 strains reviewed. Specimens where the craniofacial region was impacted by imaging artifacts, severe post-mortem damage, or large scale deformity that prevented expert identification of corresponding landmark points were excluded from asymmetry analysis.**

| Strain |
|---|
| *Acvr2b* |
| *Aebp1* |
| *Ampd1* |
| *Ano6* |
| *Arid1b* |
| *Baiap2* |
| *Bckdha* |
| *Casq2* |
| *Casr* |
| *Cbx4* |
| *Cdkn1c* |
| *Chst14* |
| *Clps* |
| *Creb3* |
| *Dcc* |
| *Fam20c* |
| *Krt5* |
| *Nhlh1* |
| *Pax7* |
| *Pdx1* |
| *Rfx6* |
| *Robo4* |
| *Slc6a5* |
| *Wnt1* |

**Table S2: Knockout strains included in analysis with no recorded structural phenotype**

| Strain | Nose | Maxillary region | Mandibular region | Eyes | Otic Placode |
|---|---|---|---|---|---|
| *Ccdc186* | 2 of 4 | 3 of 4 | 2 of 4 | 2 of 4 | 2 of 4 |
| *Acvr2a* | 2 of 4 | 2 of 4 | 0 of 4 | 2 of 4 | 1 of 4 |
| *Nhlh1* | 2 of 6 | 0 of 6 | 1 of 6 | 1 of 6 | 2 of 6 |
| *Fam20c* | 4 of 6 | 2 of 6 | 1 of 6 | 1 of 6 | 3 of 6 |

**Table S3: The number of specimens with significant asymmetry for knockout strains with the highest values of asymmetry.**

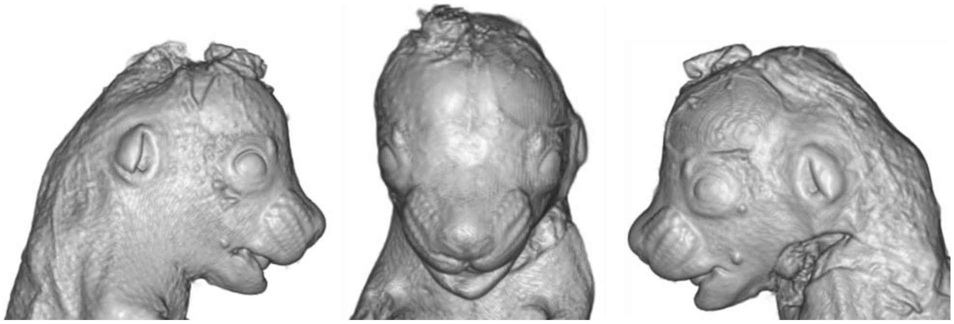

(a) *Ccdc186*

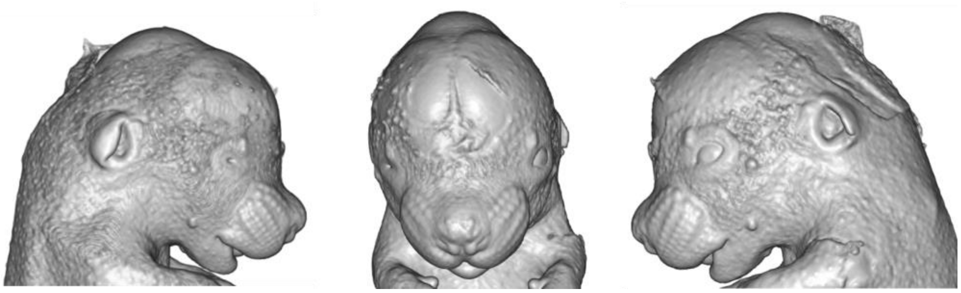

(b) *Nhlh1*

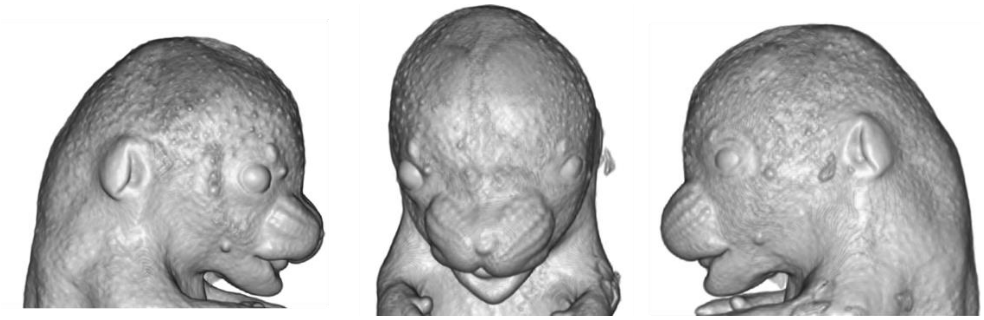

(c) *Acvr2a*

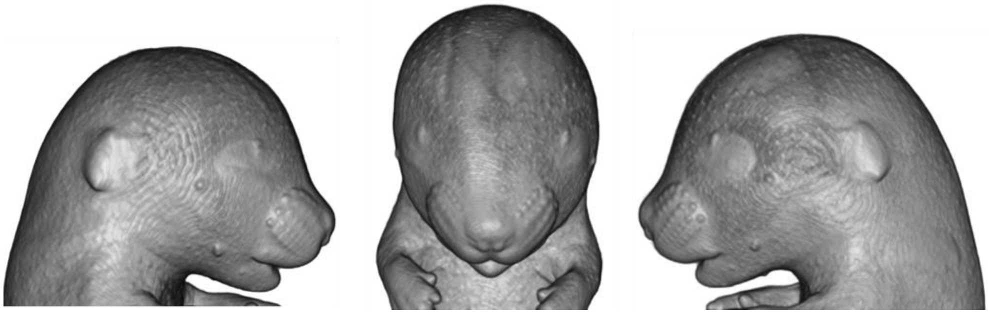

(d) *Fam20c*

**Figure S1: Right, center, and left views of the 4 specimens with significant asymmetry in the craniofacial regions: (a) *Ccdc186*, (b) *Nhlh1*, (c) *Acvr2a*, and (d) *Fam20c.***

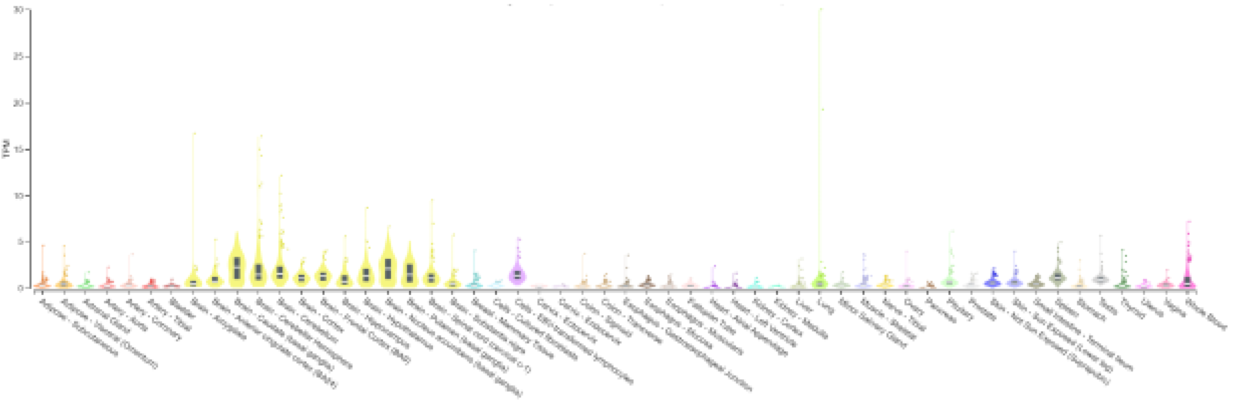

(a) NHLH1

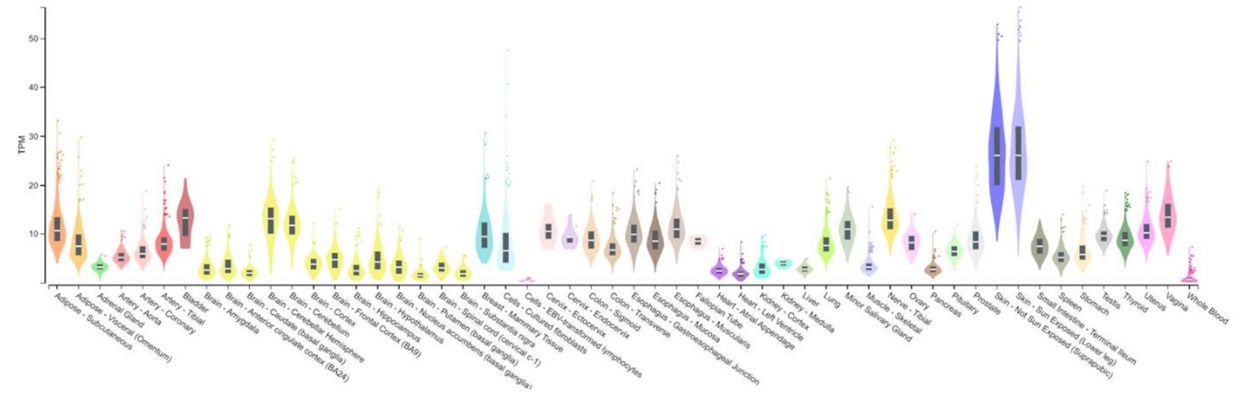


(b) ACVR2A

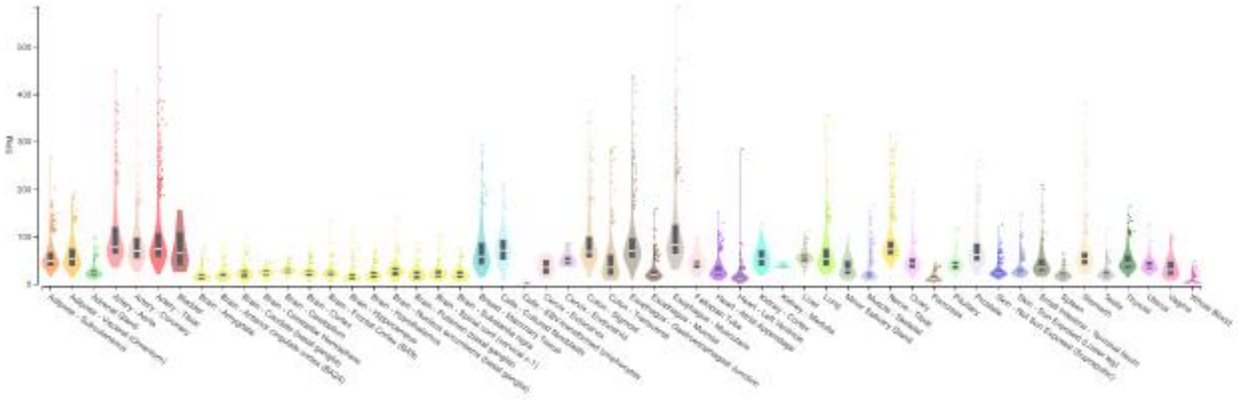

(c) FAM20C

**Figure S2: Bulk human tissue gene expression from the GTEx database for (a) NHLH1, (b) ACVR2A and (c) FAM20C, showing widespread expression of the gene across sampled tissue types in humans.**
